# The effects of age, sex, weight and breed on canid methylomes

**DOI:** 10.1101/2021.10.05.463246

**Authors:** Liudmilla Rubbi, Haoxuan Zhang, Junxi Feng, Christopher He, Patrick Kurnia, Prashansa Ratan, Aakash Tammana, Michael Thompson, Daniel Stahler, Elaine A. Ostrander, Bridgett vonHoldt, Matteo Pellegrini

**Affiliations:** Molecular, Cell and Developmental Biology, University of California Los Angeles; Yellowstone Center for Resources. Yellowstone National Park, Wyoming 82190 USA; National Human Genome Research Institute, National Institutes of Health, Bethesda MD 20892; Department of Ecology and Evolutionary Biology, Princeton University, Princeton, New Jersey 08544 USA

## Abstract

Unlike genomes, which are static throughout the lifespan of an organism, DNA methylomes are dynamic. To study these dynamics we developed quantitative models that measure the effect of multiple factors on DNA methylomes including, age, sex, weight and genetics. We conducted our study in canids, which prove to be an ideal species to assess epigenetic moderators due to their extreme variability in size and well-characterized genetic structure. We collected buccal swabs from 217 canids (207 domestic dogs and 10 gray wolves) and used targeted bisulfite sequencing to measure methylomes. We also measured genotypes at over one thousand single nucleotide polymorphisms (SNPs). We found that DNA methylomes are strongly associated with age, enabling the construction of epigenetic clocks. We also show that methylomes are strongly impacted by sex, weight and sterilization status, leading to accurate predators of these factors. Methylomes are also affected by genetics and we observe multiple associations between SNP loci and methylated CpGs. Finally, we show that several factors moderate the relationship between epigenetic ages and real ages, such as body weight, which increases epigenetic aging. In conclusion, we demonstrate that the plasticity of DNA methylomes is impacted by myriad genetic and physiological factors, and that DNA methylation biomarkers are accurate predictors of age, sex and sterilization status.

## Introduction

Domestic dogs (*Canis lupus familiaris*) and humans share a long and intertwined history, reflected by their roles as herders, hunters, guardians and companions. Domestication likely initiated 11,000-16,000 years ago (Freedman et al., 2014). Domesticated dogs inhabit the same environments as humans and are influenced by similar shared factors including pollution and disease-causing microbes (Gilmore and Greer, 2015; Sándor and Kubinyi, 2019). In addition, dogs and humans share many fundamental processes including those relating to development, disease susceptibility, behavior patterns, and aging (Sándor and Kubinyi, 2019; Wang et al., 2020). For these reasons, there has been significant interest in the study of canine genetics and, more recently, epigenetics.

Several factors affect canine lifespan. One remarkable aspect of trait variability in dogs is the extreme variation noted in body size and weight between breeds. The genetic underpinning of that variation are well established, with a small number of loci driving much of the variability (Plassais et al., 2019). Several studies have shown that body weight interacts with lifespan whereas larger breeds have shorter lifespans and generally age faster than smaller breeds (Greer et al., 2007; Kraus et al., 2013). Many breeds have, and continue to undergo strong human selection to meet breed standards set by registering bodies such the American Kennel Club (AKC). Additionally, dog lineages frequently reflect crosses between closely related individuals in an effort to maintain breed-defining phenotypes. However such practices can negatively affect breed health and impact lifespan (Yordy et al., 2020). Finally, the reproductive capability of a dog can have a significant impact on lifespan, and sterilization has been shown to increase longevity (Hoffman et al., 2013).

While there have been myriad investigations into dog genetics and their relationships to multiple traits, significantly fewer studies have addressed the relationship between canid epigenetics and traits. DNA methylation is a covalent modification, which consists in the addition of methyl groups to the C5 position of cytosine, preferentially at CpG dinucleotides. It has been implicated in the regulation of gene expression and is generally regulated by the methylation of the fourth lysine of histone H3, which preferentially occurs at promoters and enhancers (Fu et al., 2020). DNA methylation can be measured using sodium bisulfite conversion, a well-established technique that can be used to profile the status of methylcytosine across the genome at single nucleotide resolution.

While genetics are fixed throughout the lifespan of an organism, DNA methylomes are dynamic, changing with age in a tissue-specific manner. Moreover, epigenomes are impacted by environmental and physiological states of organisms. For example, body mass index (BMI) and adiposity have been shown to have observable effects on DNA methylation (Mendelson et al., 2017; Shah et al., 2015; Wahl et al., 2017). Moreover, there is a well-established relationship between methylation changes and aging, which has led to the establishment of epigenetic clocks that predict the age of an individual based on their DNA methylation profile (Thompson et al., 2017).

In this study, we measure the impact of age, sex, weight and sterilization status on canid epigenomes. We carry out these measurements across several dozen breeds, and have also included a limited number of wolves. The DNA methylation profiles are measured using targeted bisulfite sequencing of DNA samples collected from buccal swabs. This approach also allows the extraction of genetic information and the identification of single nucleotide polymorphisms from a minimally invasive tissue collection procedure. We show that all of the above traits, along with a set of associated genotypes have significant impacts on canid epigenomes. We quantify these effects by developing several models that predict age, sex, sterilization status, etc. directly from methylation profiles, and show how aging epigenetic biomarkers are in turn affected by other traits. These results demonstrate that the epigenome serves as a central molecular hub for the integration of genetic, physiological and environmental data.

## Results

### DNA methylation changes associated with age

To study age associated change in DNA methylation, we used two complementary approaches: the epigenetic clock and the epigenetic pacemaker. Epigenetic clocks have been constructed for multiple species to predict the age of an animal based on their methylation profile. These clocks are typically constructed using machine learning approaches that allow for the generation of robust age estimators. Here we used Least Angle Regression (LAR) to arrive at an age predictor that consists of a weighted sum of methylation sites. As the methylation data is high dimensional with over five thousand methylation measurements per sample, we used LAR to reduce the complexity of the search space. To avoid overfitting, we used leave-one-out cross-validation to build a separate model for each sample that is trained on all the remaining data once that sample has been excluded. We found that the resulting model is able to accurately predict the age of dogs to within an average error of approximately ten months (Fig 1A). Based on previous work, we also hypothesized that DNA methylation changes may not be linear with time, rather, methylation profiles may change more rapidly earlier in life. To account for such non-linearities, we trained a model for the square root of age which approximates a trend in which methylation changes more slowly with increasing age. We observe that the model trained on the square root of age slightly outperforms the linear model, with a median absolute error of eight months (Fig 1B).

**Fig 1.**
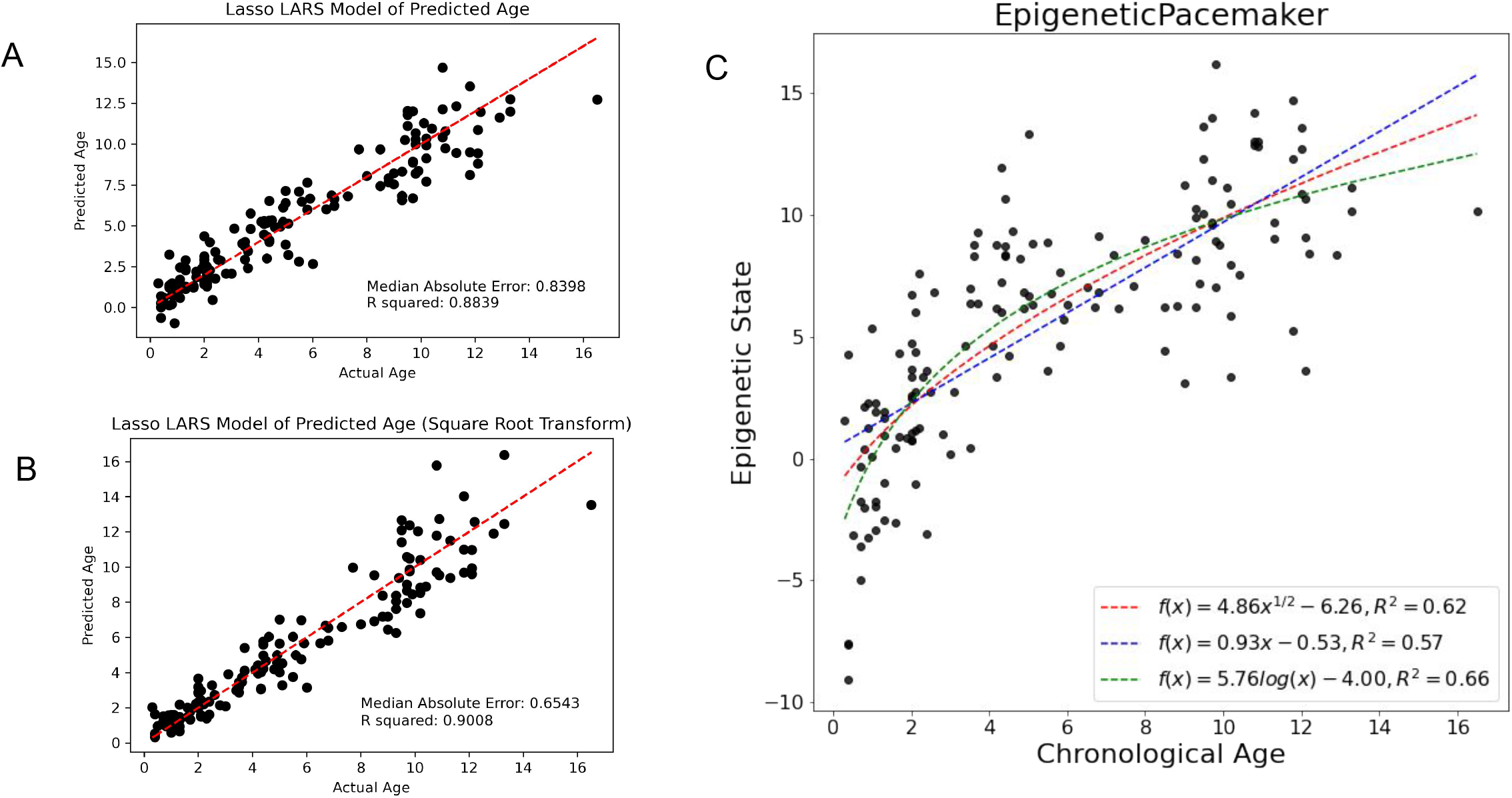
Epigenetic age and state of dogs. (A) Models generated using least angle regression (LARS). (B) LARS model of the square root of age. (C) Epigenetic Pacemaker was also used to predict epigenetic states, and the trend line was fit using three functions - linear, logarithmic, and square root.

While epigenetic clocks are useful predictors of age, as the examples above demonstrate they require *a priori* assumptions about the functional form of age associated methylation changes. Our previous analysis suggests that the square root function may more accurately characterize age associated DNA methylation changes. However, it is of interest to identify this functional form in an unbiased fashion. To this end, we have previously developed the epigenetic pacemaker (Farrell et al., 2020). This approach is distinct from an epigenetic clock, in that it attempts to model the time dependent changes in DNA methylation directly, rather than predict an age based on methylation levels. The underlying model is described by the following equation:

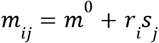

Here *m*_*ij*_ represents the methylation level of position *i* in individual *j, m*^0^ represents the methylation level at birth,*r*_*j*_the rate of change of methylation with respect to an underlying epigenetic state *s*_*j*_. The epigenetic state of each individual represents a position in the epigenetic trajectory of its lifespan. To solve the model, we use an expectation minimization approach in which we recursively estimate the epigenetic state, then *m*^0^ and *r*_*j*_, and repeat the process until the model converges and minimizes the difference between the observed and predicted methylation values in our dataset. The advantage of this approach is that the relationship between the epigenetic state and age is not constrained and is optimized by the model. We previously found that a logarithmic function offers the best fit for age-epigenetic associations in humans when the dataset spans a very broad age range (Snir et al., 2019). Here, we find a similar result, with the logarithmic trend line best describing the relationship between the epigenetic state of dogs and their age (Fig 1C). This result suggests that DNA methylation is constantly decreasing as an animal ages, and the rate of change is approximately inversely proportional to the age of the dog.

### Effect of sex, sterilization and weight on DNA methylation

While many studies demonstrate a relationship between DNA methylation and age, the effects of other traits on methylation have been less well characterized. We first asked whether the methylome can distinguish male from female dogs. We trained a lasso logistic regression model, using leave one out cross validation, to test if methylatioh profiles accurately predict the sex of dogs. Not surprisingly, we found that such a model had nearly perfect performance and could accurately classify all but one dog (Fig 2A). We also asked whether certain sites in our methylome had a stronger influence on these models and found that the most informative sites were on the X chromosome (Supp Fig 1). This result supports the notion that the male and female X chromosomes are differentially methylated, likely due to chromosome inactivation of one of the X chromosomes in females (Duncan et al., 2018). Indeed, this inactivation is associated with hypermethylation of the inactive X chromosome. Thus, the single male X chromosome has many differentially methylated loci when compared to the combined active and inactive female X chromosomes. This finding is concordant with a past study of dog and gray wolf methylation profiles across their genomes (Koch et al., 2016).

**Fig 2.**
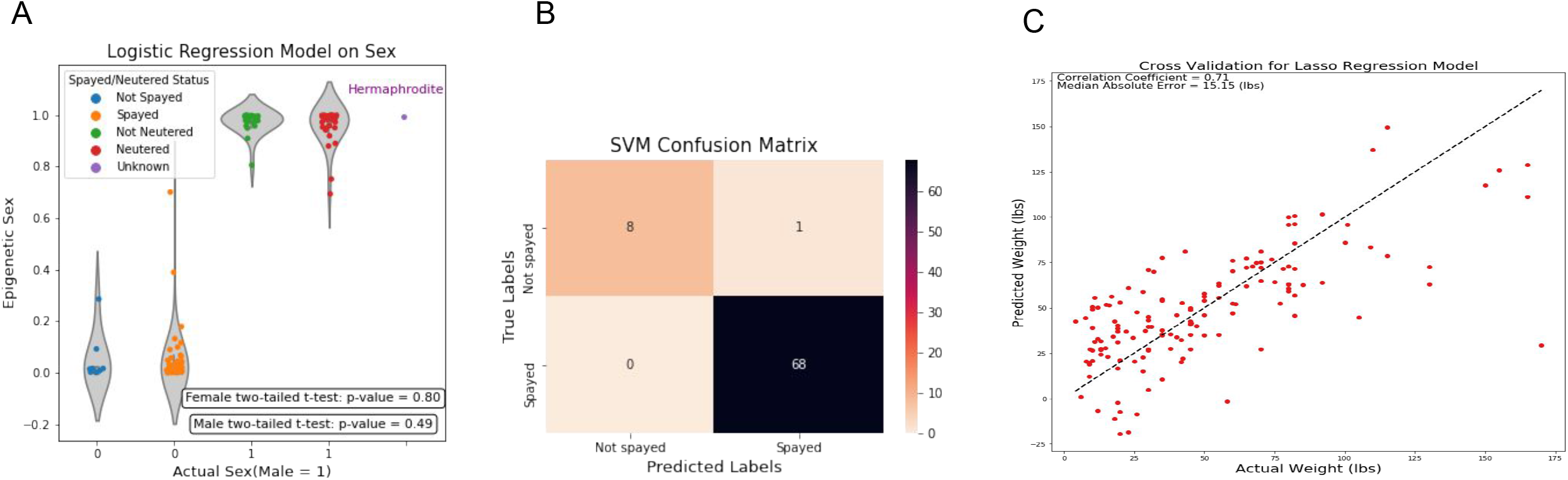
Epigenetics of sex, weight and spayed status. (A) Predicted sex values (labeled as Epigenetic Sex on the y-axis) were colored based on the sample’s sterilization status into five categories: blue for not spayed, orange for spayed, green for not neutered, red for neutered, and purple for missing values. A two-tailed t-test was performed separately for each sex, on the predicted values of fixed and intact samples. The hermaphrodite (intersex) wolf was not included in the t-tests. (B) The confusion matrix displays the number of correctly and incorrectly predicted spayed status for the female samples during the LOOCV process. The rows are the true sterilization status, while the columns are the predicted sterilization status by the SVM model. (C) The lasso model from sklearn is used to predict the weight of dogs.

As our collection of dogs contained both sterilized (i.e. spayed or neutered) and intact dogs, we tested whether the epigenetic sex prediction reported above could differentially distinguish sterilization status between males and females. However we found no statistically significant differences between either male or female sterilized versus intact dogs (Fig 1A). This finding suggests that differential methylation at X chromosomes sites are not strongly influenced by hormonal levels that differ between sterilized and intact dogs However, we note that the variance of the sex prediction for sterilized dogs was significantly higher than for intact dogs (P < 1e-4 for a one way F test), suggesting that there may be an increase in the variability of the methylation at these sites in response to spaying or neutering.

Interestingly, one of the canids sequenced was an intersex gray wolf. Intersexuality occurs in domestic dogs but has also been described in wild gray wolves (Kang et al., 2012; Poth et al., 2010). This individual reflected a rare discovery sampled via capture and handling efforts in the Yellowstone region, with over 1,200 individuals handled during the last 25 years. Usually, we use the naming convention of a “M” or “F” behind the field ID number to denote sex, but this individual wolf recieved a “U” for unknown sex. When this wolf was first captured and handled in January 2018, it’s age was estimated at 5-6 years. Although there was limited observational data for this wolf, there were a few documented facts. First, it traveled with a male packmate 1117M and they seemed to act like a bonded male/female pair. Features of the intersex wolf included a vulva at the location where the preputial orifice would normally be located in a male and, it had an os penis, but no functional penis. The urinary system functioned like a female except that the vulva was on the belly rather than under the anus. It had one descended testicle in the scrotum while the other was retained in the abdomen. It also had a partially formed uterus, with a uterine horn on the right, half a uterine horn on the left, but no ovaries. Interestingly, the epigenetic sex prediction clearly classified this individual as a male. This result supports the notion that sex-specific hormonal and genitalia differences seem to have limited effects on epigenetic sex predictions.

Although the effects of sterilization on the epigenetic sex prediction seemed to be muted, we nonethelss asked if we could use the DNA methylation profiles to predict a dog’s sterilization status. To this end we trained a Support Vector Machine (SVM) using leave-one-out cross-validation. We found that while the SVM was unable to accurately predict the state of the males, it could very accurately predict the sterilization status of females (Fig 1B). This result may indicate that the female methylome is more susceptible to the hormonal changes induced by sterilization than a male methylome.

Lastly, we asked if the canid methylome is impacted by the weight of a dog. We used a similar approach as for age above, and constructed a penalized regression model for weight using leave-one-out cross-validation. We found that the weight of a dog could not be as accurately predicted as its age. The model had a correlation coefficient of 0.7 and a median absolute error of 15 pounds. We noted that a few of the outliers were heavy breeds (> 150 lbs?) that are predicted to be of low weight during their first year of life, suggesting that the epigenetic weight of a dog may reach its final level only once the dog has reached adulthood. To test this observation, we trained a model on the weight times the square root of the dog’s age and found that it outperformed the previous model, supporting our notion that the effect of weight on the methylome may depend on age (Supplementary Fig 2).

### Dog genetics and association with weight

Due to the nature of bisulfite converted DNA, some genotype information is lost at each cytosine, while it is preserved at other nucleotides. To circumvent this issue, we used a bisulfite-aware genotyping approach to extract the genotypes of all non-cytosine bases in the capture regions, which were then further refined to identify variable regions and SNPs. Using this approach, we identified approximately one thousand five hundred SNPs, or about one SNP for every three probes (see Methods).

We used the SNP genotype data and constructed a phylogenetic tree of the samples (Fig 3A). We first noted that dogs of the same breed tended to group together, supporting our ability to correctly reconstruct breed relationships. Moreover, as expected, we observed that the wolves were an outgroup to the dogs. Among the dogs, we found that the chow-chow breed was the first diverged group from the gray wolves, followed by the saluki. These relationships are supported by previously published breed relationships based on SNP arrays (Parker et al., 2017). The remaining breeds resulted from more recent breeding and formed one large clade of several subclusters that represent functional breed groupings or history of breed formation.

**Fig 3.**
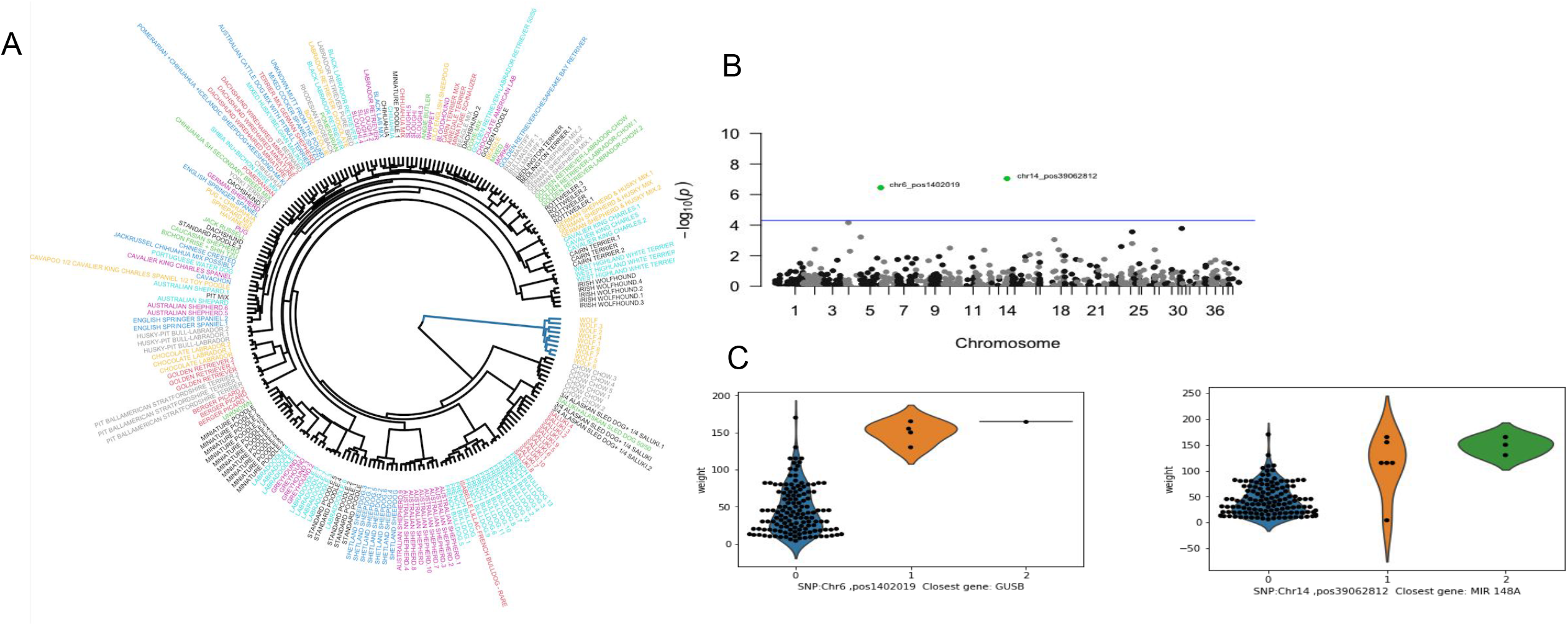
Phylogenetic tree and weight GWAS. (A) Different breeds of dogs are shown on a hierarchical clustering tree based on their genotypes and each breed is assigned with a unique color. (B) A genome-wide association analysis is performed on the weight of dogs, and the results are shown in a Manhattan plot. Significant SNPs are marked in green, along with their chromosome number and position, and the weight of dogs with each genotype is shown as a violin plot.

Among the traits measured only weight was considered to have a genetic component in our SNP data, as age, sex and sterilization status should not depend on genetics. We therefore conducted a genome wide association study (GWAS) to identify SNPs that are associated with weight (Fig 3B). Two SNPs reached statistical significance after correction for multiple testing: chromosome CFA6:1402,019 and chromosome CFA14: 39,062,812 (Fig 3C). The closest genes to these positions are GUSB and MIR-148A, respectively. GUSB is a beta-glucuronidase, an enzyme that breaks down glucuronic acid. Deficiency of beta-glucuronidase in dogs results in mucopolysaccharidosis VII (MPS VII), also known as Sly Syndrome, which is related to the physiological development of dogs (Haskins et al., 1984; Ponder et al., 2002). MIR-148A is a microRNA gene and has been shown to serve as a biomarker of obesity in humans (Shi et al., 2015). Neither of these SNPs has been associated with dog weight in previous GWAS studies, but this may be due to the poor overlap between our limited number of target loci and the SNP panels on the existing arrays.

### Interactions between genotypes and DNA methylation

We next carried out an association study to identify significant correlations between genotypes and DNA methylation levels (Fig 4A). We identified only one significant *cis* association between the SNP at position CFA25:25,099,323 and the methylation of the probe at CFA25:25,099,291 (adjP<10^−12^; Fig 4B). By contrast, we found four significant *trans* associations at a significance level of adjP<10^−4^ (Fig 4C). Of particular interest was the observation that only two SNPs were involved with the four significant associations, and that one of these SNPs marks a position that is a hotspot for DNA methylation regulation. The hotspot with the three associations was proximal to the gene encoding HMG20A. Although we do not have eQTL data, it is reasonable to assume that this SNP may influence the expression of this gene and that HMG20A in turn modulates DNA methylation levels via *trans*-regulation. HMG20A is a high mobility group (HMG) domain containing protein homologous to HMG20B, a core subunit of the Lys-specific demethylase 1/REST corepressor 1 (LSD1-CoREST) histone demethylase complex (Rivero et al., 2015, p. 20). The LSD1 complex has been associated with the demethylation of H3K4 (Kim et al., 2020). Since *de novo* DNA methyltransferases are inhibited by H3K4me3 (Fu et al., 2020), it is reasonable to assume that the regulation of the expression of HMG20A could affect the demethylation of H3K4 loci in *trans* and hence their DNA methylation levels.

**Fig 4.**
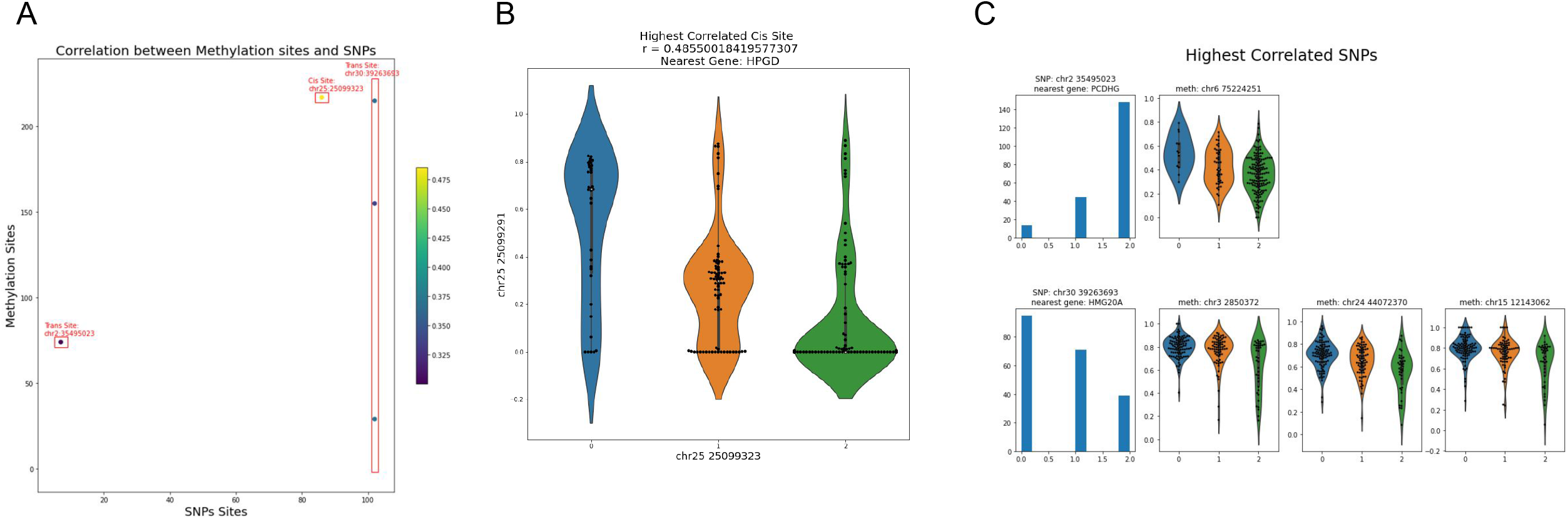
Association of SNPs and methylation values in cis and trans. ((A) We identified 5 significantly correlated pairs of SNP and methylation sites using pearson correlation and the Benjamini-Hochberg procedure with a false discovery rate of 0.1, only one of which is a cis pair. One SNP, at location chr30:39263693 is significantly correlated with three different methylation sites, all of which are trans. (B) Violin plot of genotype vs methylation values for the significant cis pair, which had a pearson correlation coefficient of 0.485. The nearest gene to this cis pair is HPGD. (C) Left-most column: histograms of genotypes across all samples for the two SNPs where significant trans correlations were found. Right 3 columns: violin plots of genotype vs methylation for the four significant trans correlations. Each row corresponds with one of the two SNPs that were found to have significant trans correlations..

### The effects of weight on epigenetic aging

As shown above, variation in canid methylomes are associated with aging, sex, weight, sterilization and genotypes. We next sought to determine whether the association between actual age and predicted age (or state) is moderated by these factors. To accomplish this, we generated linear models for both the epigenetic age and the epigenetic state using the other factors as moderators. We measured whether, for example, the epigenetic age could be modeled using the age of a dog plus the interaction between age and weight. We infer that any term other than age itself found to be significant in such models moderates the association between actual and epigenetic age. Using this approach we tested several moderators including weight, sex, sterilization, heterozygosity and runs of homozygosity.

Of all of the factors we tested only weight significantly moderated the effect of age on epigenetic aging (Fig 5A). The effect of weight on epigenetic ages is illustrated in Fig 5B by dichotomizing the dog cohort into two groups based on weight and showing that they have different slopes, with the heavier dogs having the higher slope, or accelerated epigenetic aging. It is well documented that the lifespan of heavier breeds is shorter than small breeds dogs. This could be due to higher mortality rates associated with certain diseases. However, our result suggests that even at the molecular level, the methylation patterns of larger breeds show accelerated aging compared to smaller breeds, and that at 12 years heavier dogs have an epigenetic age acceleration of about two years compared to smaller dogs. Thus it appears that the weight of dogs has a direct impact on the rate of epigenetic changes.

**Fig 5.**
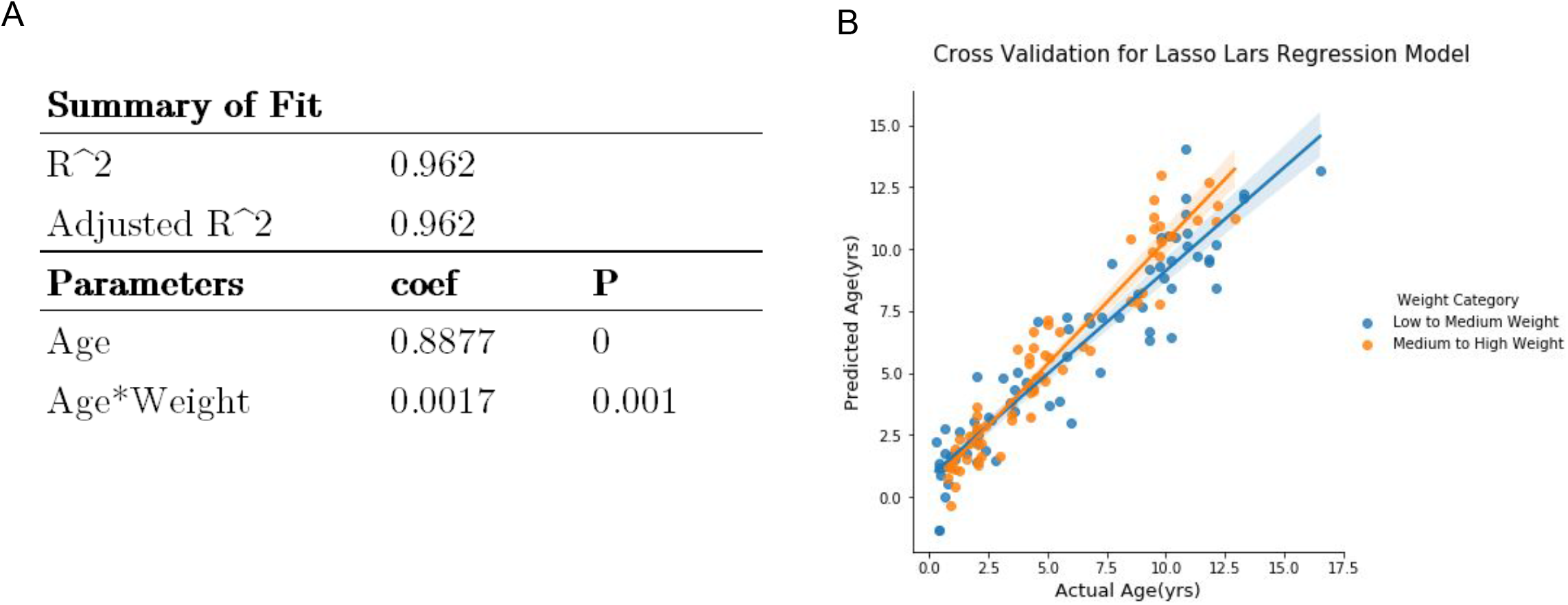
Weight moderates epigenetic age. (A) The predicted age can be expressed as a linear combination of age and age times weight with both terms being significant. (B) Weight values were sorted into the two categories, low-to-medium weight (blue) and medium-to-high weight (orange), by using 72 lbs as the cutoff and the slopes of epigenetic age vs. actual age for the two weight groups are shown,

We conducted a similar analysis by modeling the factors that moderate the epigenetic state, rather than the epigenetic age. As described above, the epigenetic state is an estimate of the position of a dog along a time-dependent trajectory of DNA methylation that represents the entire lifespan of a dog. Unlike the epigenetic age which is a model that is optimized to predict the age of a dog, the epigenetic state is optimized to minimize the error in the predicted methylation levels of a limited number of sites that manifest age-associated DNA methylation level changes. In previous studies we have observed that the epigenetic state is more impacted by external and internal factors than the epigenetic age (Pinho et al., 2021). Consistent with this observation we find that the linear model of the epigenetic state has multiple significant genetic moderators including heterozygosity (see Methods) as well as two principal components of the genotypes (Fig 6A). Since we could not include individual SNPs in this analysis due to lack of statistical power to test over one thousand loci, we used principal component analysis to reduce the dimensionality of our genotype matrix. Using a greedy algorithm to find significant moderators of epigenetic states we identified two principal components principal components 5 and 7 (Supplementary Figure 3). Although it is often difficult to interpret these components, we did observe that component 7 was significantly anticorrelated with weight, and thus may be capturing weight dependent genetic effects (Supplementary Fig 4). The effects of these genetic moderators of the epigenetic state are shown in Fig 6B. These results further illustrate the interactions of the methylome, as characterized by the epigenetic state, with genetic factors.

**Fig 5.**
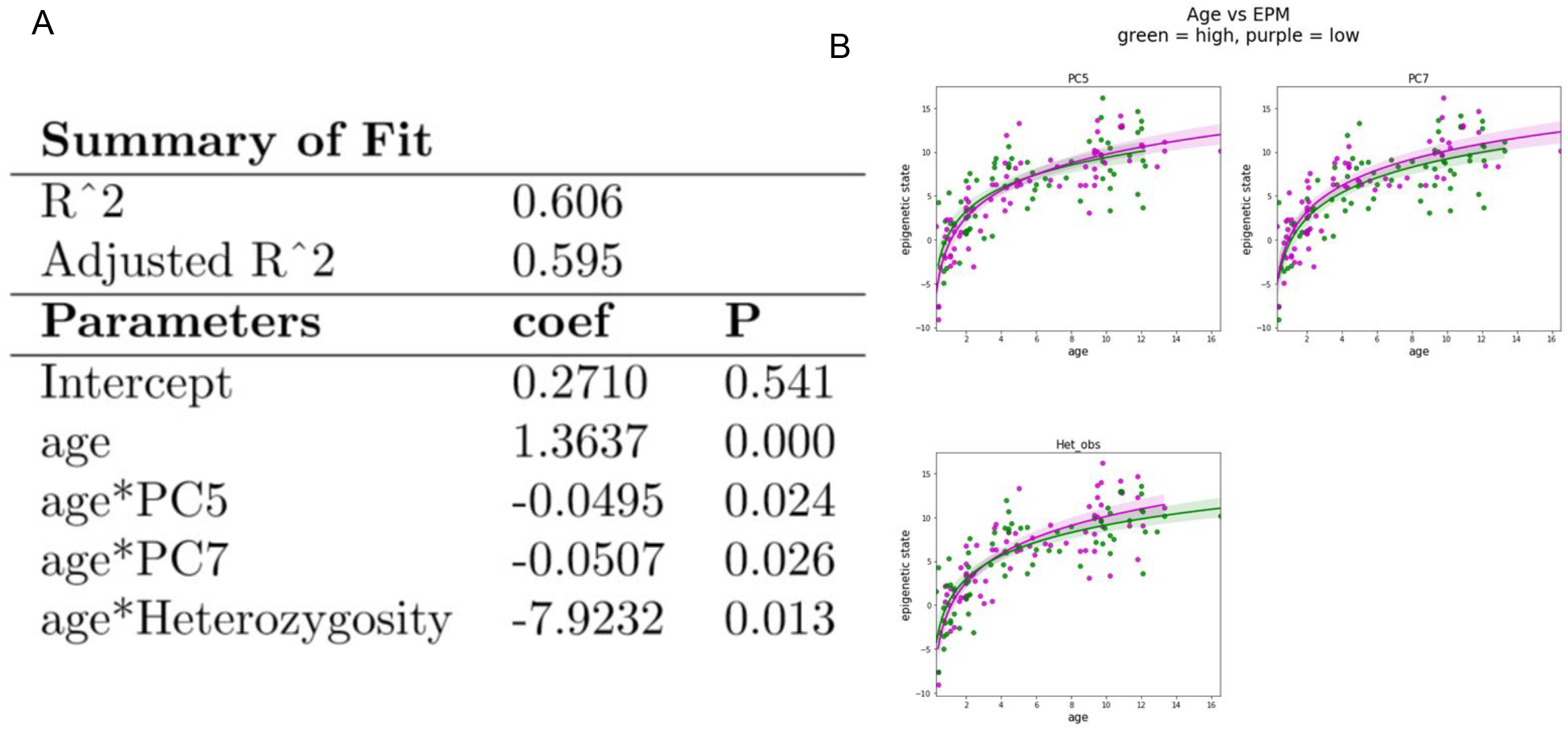
Genetics impacts epigenetic states. (A) Linear model of epigenetic states including heterozygosity and principal components 5 and 7, all of which had significant p-values. Significant traits were selected from weight, sex, sterilization status, heterozygosity, and the top 10 principal components using a greedy algorithm. (B) Age vs epigenetic state plots split by value of the 3 significant traits: PC5, PC7, and heterozygosity. The purple points are samples with values higher than the median and the green points are samples with values lower than the median.

## Discussion

Unlike the four letter sequence of germline DNA that is largely invariant across both spatial and temporal dimensions in an organism, the five letter epigenetic code that includes methylcytosine shows great variability between cell types and during the lifespan of an organism. Changes in the methylome are driven by myriad signals that include environmental and physiological changes along with developmental programs that are most influential early in life. Thus, unlike the genome, the methylome accumulates multiple changes during a lifespan and can be viewed as a historical record of an organism. Deciphering the factors that most strongly impact this record remains one of the central goals of epigenomics.

Most studies on methylomes to date have been conducted on humans and have described associations between methylation, age and disease. Non-human studies are still more limited but have many advantages over human studies as they can probe phenotypes not easily accessible in humans. To this end, dogs are a valuable model organism for reconstructing the impact of environment and physiology on methylomes. The lifespan of dog breeds, which ranges from about a low of eight years through an average of 10-13 to a high of 18 is intermediate between that of humans and mice. Therefore aging timescales in dogs are much more similar to those of humans. Moreover, breeds show remarkable phenotypic plasticity compared to humans and are therefore ideal for studying the interactions between body size factors and DNA methylation. For example, the weight range for dog breeds varies by 40-fold, making it much larger than the range observed in human populations. Genetic variation between dog breeds and wolves likely represents approximately 20,000years of evolution and domestication. While many modern dog breeds are genetically homogeneous, domestication followed by human driven selection has resulted in variants strongly associated with specific traits.

In this study we set out to measure the impact of age, sex, weight, sterilization and genetics, on the methylome in an attempt to characterize factors that influence the plasticity of the methylomes in canids. We profiled dog DNA from buccal swabs, as this tissue is easily accessible via non-invasive methods, and also included DNA from the blood of a small number of wolves in our study. Because of the ease of collection of buccal swabs our results should be easily validated and extended in future studies and possibly leading to DNA tests that can be offered to the public at large.

To profile DNA methylation patterns, we used a targeted bisulfite sequencing approach. Despite rapid advances in the efficiency of DNA sequencing, it is still costly to generate whole genome bisulfite sequences for hundreds of samples. To reduce the sequencing depth needed, we designed a panel of probes that capture a few thousand loci in the genome. These were selected based on previous studies to identify loci whose methylation changes with age (Thompson et al., 2017). We also included loci that are hyper conserved across mammals, as these have been shown to contain regulatory regions that are differentially methylated during development and between cell types (Colwell et al., 2018). Although the panel we developed captures a very small faction of the CpG sites in the genome, they are strongly enriched for sites with variable methylation and are therefore suited for the study of factors that impact methylomes.

In support of previous studies that investigated the changes in DNA methylation with age, we found that the age of a dog has a significant impact on its methylome. We were able to study the relationship between methylomes and aging using two different approaches. In the first we built a model to predict the age of a dog from its methylation patterns and found that we could do so with an accuracy of approximately eight months. We also used a second approach called the Epigenetic Pacemaker, to directly model DNA methylation profiles in a time dependent fashion. This approach allows us to identify the epigenetic state of each individual along a trajectory that summarizes DNA methylation changes across a lifespan. With this approach we were able to confirm that DNA methylation changes are more rapid early in life and slow down as a dog ages, a trend that can be accurately approximated using a logarithmic function. Similar trends have also been observed in human studies (Snir et al., 2019), and are consistent with the notion that DNA methylation changes rapidly during embryogenesis and childhood as the body shape of an organism is changing, and then slows down, but never stops changing as one ages.

We also find that DNA methylation profiles differs between males and females, and can be used to develop accurate predictors of sex. Similar sex predictors have also been constructed for humans and have proven to be highly accurate (Wang et al., 2021). Not surprisingly, most of the sites that contribute strongly to our sex prediction model are found on the sex chromosome. Females have two X chromosomes, one of which is epigenetically inactivated. The inactive sex chromosme has a distinct DNA methylation profile from the active one (Duncan et al., 2018). Therefore, it is not surprising that the combination of the two female X chromosomes have slightly different DNA methylation patterns than the single male X chromosome. We were also able to test whether sex prediction using our DNA methylation based classifier were influenced by sterilization. We found that the accuracy of predictions was not significantly impacted, but we did find a significant difference in the variance of the prediction for intact and sterilized animals, suggesting that sterilization does increase the variance of methylation in the sex chromosomes, even though the underlying patterns are still distinct between males and females. In support of this conclusion, we were able to train a classifier to predict sterilization, and found that we could accurately predict whether a dog was spayed, but not whether it was neutered. Thus, the hormonal changes induced by sterilization of females have significant impacts on the epigenome, even though sites that are differentially methylated between males and females are largely retained. The fact that we found more significant impacts from spaying than neutering might indicate that changes in female hormone state have a larger impact on the epigenome than changes in male hormone state, as has been observed in humans (Thompson et al., 2018). We also examined the curious case of an intersex wolf. Despite this animal having both male and female genitalia, the epigenetic sex prediction model classified this individual as a male, further reinforcing the observation that epigenetic sex states are not strongly affected by hormonal changes that impact the development of genitalia.

We also find that the methylome of dogs is significantly influenced by their weight. By constructing an epigenetic predictor of dog weight, we are able to show that although not as accurately predicted as age, the epigenetic weight was very significantly associated with the weight, generating predictions with a median error of about fifteen pounds. Of interest was the observation that a few of the outliers in our model, whose epigenetic weight was significantly underpredicted, were less than one year old. As it is well known that the adult weight of a dog is not reached until after the age of one, this suggests that the effect of the adult weight on the epigenome may continue to change, even after the weight has stabilized.

Since both the weight and the age of the dog impact the epigenome, we also asked whether these two factors interact to moderate the association between real and epigenetic age. It is well established that the lifespan of heavier dogs tends to be shorter than smaller dogs (Michell, 1999). The lifespan of the smallest compared to the largest breeds can differ by a factor of two. There are many possible explanations for the effect of accelerated aging in heavier breeds, and previous studies had suggested that there may be a corresponding increase in epigenetic aging with dog weight (Thompson et al., 2017). In our cohort we observed that there was a significant association between epigenetic aging rates and weight. As the accelerated aging of large breeds is manifest in DNA methylation patterns, it appears to be an intrinsic molecular property of dog aging rates. We still do not know why this is the case, but speculate that accelerated developmental programs that lead to larger body sizes may continue to influence the rate of epigenetic changes even into adulthood, thus leading to faster epigenetic aging rates throughout a lifespan.

Although the epigenetic clock was only significantly affected by a dog’s age and weight, the epigenetic state of each dog proved to be influenced by multiple genetic factors. Since the epigenetic state is not trained to predict age, but rather attempts to model time dependent methylation changes, it is not surprising that this approach is more sensitive to genetic influences on the epigenome than an epigenetic clock. We found that two of the principal components that were derived from dog genotypes significantly interacted with age to modify the epigenetic state. One of these components, PC7, was also significantly associated with weight, and thus may capture some of the genetic variation that is related to breed size. Thus the epigenetic pacemaker paradigm was able to capture multiple factors that interact with age to impact the epigenetic state of a dog, further demonstrating the complex link between genetics, and epigenetics.

In conclusion, we have shown that the epigenome of a dog is affected by myriad factors that can be identified using quantitative models of the age or of the methylation landscape itself. As the targeted bisulfite sequencing methodology we have developed allows us to extract both epigenetic and genetic information, this approach can be applied extensively across varied species to further elaborate interactions between epigenetics, genetics, physiology and environment. The quantitative modeling of these interactions will bring us closer to the ability to maximally extract information from the epigenome, so that we can complement static genetic data with dynamic epigenetic profiles to more fully characterize the phenotype of an organism from their DNA.

## Materials and Methods

### Canid Samples

A total of 217 DNA samples were collected from three different sources: 10 gray wolves were part of the Yellowstone National Park cohort. A total of 207 dogs were sampled from two cohorts. The first set of 151 dogs were sampled using buccal swabs purchased from DNA Genotek (Performagene PG-100 device). For each saliva sample the following details were collected: date of birth (accuracy to the month and year of the birth), breed, weight, sex, and spayed/neutered status. Samples from these dogs were collected with the owners’ signed consents. The samples were collected from dog parks and breeders and online volunteers who mailed in samples.

The remaining 56 samples provided by the National Human Genome Research Institute (NHGR) of the National Institutes of Health were collected in accordance with all NHGRI Care and Use Protocols, with owner’s signed consent. Only samples of known breed and usually with AKC number were sampled for this study. Dogs were unrelated at the grandparent level. For each sample, age of the dog at collection, sex, breed, and was provided. A range of sizes and breed types (terriers, working dogs, spitz breed, etc.) were provided. Individual weights were not available on dog samples, hence breed standard weights were used based on American Kennel Club breed descriptions. Cheek swab samples were collected using PERFORMAgene kit (PG-100) from DNAGenotek and DNA was isolated according to the manufacturer instructions. Samples were stored at −80 degrees C. Some samples were provided by collaborating institutions.

We also analyzed genomic DNA from 10 gray wolves (*Canis lupus*) from Yellowstone National Park. This population was founded in 1995 and 1996 when 41 individuals were reintroduced and has been heavily monitored since through annual capture/release seasons, daily telemetry, and direct observations. Individual DNA was extracted from whole blood with the decimal age and sex of the individual known at the time of capture. All wolves were handled in accordance with recommendations from the American Society of Mammalogists (Sikes 2016) and approved by the National Park Service’s Institutional Animal Care and Use Committee. This population is genetically pedigreed and has been extensively studied with respect to social structure, reproduction, behavior, disease, and genomics (e.g. Almberg et al., 2009; vonHoldt et al., 2020; Cassidy et al., 2017; Cubaynes et al., 2014; Stahler et al., 2013; vonHoldt et al., 2010; vonHoldt et al., 2008).

All canid characteristics are described in Supplementary table 1.

### Targeted Bisulfite sequencing

DNA was extracted from the buccal swabs using the vendor supplied protocol. Buccal swabs were incubated overnight at 50C degrees before DNA extraction.

We applied targeted bisulfite sequencing (TBS-seq) to characterize the methylomes of the 217 samples. The protocol is described in detail in (Morselli et al., 2021). Briefly, 500 ng of extracted DNA were used for TBS-seq library preparation. Fragmented DNA was subject to end repair, dA-tailing and adapter ligation using the NEBNext Ultra II Library prep kit using custom pre-methylated adapters (IDT). Pools of 16 purified libraries were hybridized to the biotinylated probes according to the manufacturer’s protocol. Captured DNA was treated with bisulfite prior to PCR amplification using KAPA HiFi Uracil+(Roche) with the following conditions: 2 min at 98°C; 14 cycles of (98°C for 20 sec; 60°C for 30 sec; 72°C for 30 sec); 72°C for 5 minutes; hold at 4°C. Library QC was performed using the High-Sensitivity D1000 Assay on a 2200 Agilent TapeStation. Pools of 96 libraries were sequenced on a NovaSeq6000 (S1 lane) as paired-end 150 bases.

The probes used in the capture were designed using two criteria. We identified 2223 regions whose methylation was highly associated with age based on a previous study of canid methylomes using Reduced Representation Bisulfite Sequencing (Thompson et al., 2017). The remaining 3573 probes were obtained from ultraconserved regions across mammals (Faircloth et al., 2012). The sequence of the probes used in this study can be found in Supplementary Table 2.

### Data Processing

Demultiplexed Fastq files were subject to adapter removal using cutadapt (v2.10) (Martin 2011) and aligned to the GRCh38 genome using BSBolt Align (v1.3.0) (Farrell et al. 2021). PCR duplicates were removed using samtools markdup function (samtools version 1.9) (Li et al. 2009) before calling methylation using BSBolt CallMethylation function. To generate the DNA methylation values for each probe region, we generated a weighted average of the methylation of CpGs within 100 bases of the probe center, where the weights were determined based on sequence coverage. The methylation levels of each probe across each dog can be found in Supplementary Table 3.

To generate genotype calls from the BAM files we first used the samtool pileup command to identify the aligned bases associated with each position within a probe. Due to bisulfite conversion, we ignored all positions that contained either cytosines or thymines and only retained positions that contained adenines and guanines. We then identified the associated genotype (either AA, AG or GG) based on the counts of each of the two bases, ignoring sites with coverage less than ten, and making heterozygous calls only for sites where the minor allele frequency was above ten percent.

### Data Analysis

#### Sex Classifier

The machine learning model for predicting sex was trained using nested cross validation on the logistic regression model from scikit-learn. After removing samples with missing sex values and those that have high proportion of missing methylation values, there were 140 wolf and dog samples used for training. Methylation data were filtered so that sites with low coverage were removed, which left a total of 2420 sites. The hyperparameter tuning loop uses the LogisticRegressionCV function with 10-fold cross validation. The outer loop uses a leave-one-out cross-validation(LOOCV) method to predict the probability of each sample belonging to the two classes of sex when it’s in the test set, using the “predict_proba” function implemented in scikit-learn’s logistic regression model.

#### Sterilization Classifier

To predict the sterilization status of the samples(separately for male and female samples), several models from scikit-learn were evaluated using LOOCV on 141 dog samples, such as ridge classifier, logistic regression, random forest classifier, and SVM. Methylation data were adjusted by removing sites with low-read, leaving a total of 2420 sites. For the male samples, the models performed poorly in predicting the intact samples. The best performance was achieved with the SVM model for females. A total of 77 female samples were used during the training process, with 9 intact and 68 spayed samples. To handle this imbalanced class distribution, the class_weight parameter in the SVM model was set to “balanced.” A parameter grid was defined to tune the hyperparameters C and gamma. The values of C to be searched were 0.01, 0.1, 1, 10, and 100. The values of gamma to be searched were 0.001, 0.01, 0.1, and 1. The kernel parameter was set to the default “rbf” kernel. Nested inside the leave-one-out cross-validation loop, the GridSearchCV function from scikit-learn was called to find the best parameter combination from the parameter grid defined above using a 10-fold cross-validation. The predicted class values were then retrieved and shown using a confusion matrix.

#### Epigenetic age prediction

Prediction of dog ages was generated using the Lasso LARS model from the scikit-learn machine learning package in Python (sklearn.linear_model.LassoLars). The methylation matrix was used as the predictor variable for the target variable of age, with and without square root transformation of age. Thirteen dogs were removed from the dataset due to high proportions of missing methylation values. The methylation sites were then filtered so that no missing values are retained. The matrix for generating predictions used 147 dogs from the two cohorts, excluding the wolves, and a total of 2358 methylation sites. Sklearn.model_selection.LeaveOneOut was used to generate individual predictions; each predicted dog represented the testing set and the training set were all other dogs.

#### Epigenetic Pacemaker

The Python package EpigeneticPacemaker (EpigeneticPacemaker.EpigeneticPacemaker) was used to generate predictions for each dog’s epigenetic state. The predictions relied on selecting methylation sites that were highly correlated with age, determined using the Pearson correlation coefficient. The minimum correlation threshold was set to be 0.6 which returned 124 sites. These sites were used to generate a plot of each dog’s predicted epigenetic state against their true age. Points were fitted to three possible trend lines - linear, logarithmic and square root - using Scipy (scipy.optimize.curve_fit).

#### Moderation analysis

To identify factors that moderate the relationships between actual and predicted age we used linear models and computed the P value of each term. Multiple factors were tested in the Lasso LARS predictions of the epigenetic age model. We used 147 dogs to train the model. The moderators we tested were weight, sex (1 = female, 0 = male), sterilization (1 = sterilized, 0 = not sterilized), heterozygosity and inbreeding. The significance of each term was analyzed using the Python package Statsmodels (statsmodels.regression.linear_model.OLS), with the matrix of factors input as the independent variables and the epigenetic age predictions as the dependent variable. The same approach was used to identify moderators of the epigenetic state.

#### Phylogenetic Tree

To generate the phylogenetic tree, the genotype matrix was first imputed using the R package ‘knncatImpute’, which applies weighted k nearest neighbor. Next, the genetic distance was calculated by the method Nei Distance with the R package ‘NAM(Nested Association Mapping)’. Groups are generated using agglomerative hierarchical clustering analysis, using the ward.D method. Finally, the phylogenetic tree was drawn with the R packages ‘dendextend’ and ‘circlize’.

#### GWAS

Genome Wide Association Analysis was performed using the R package ‘statgenGWAS’. The package implements the Generalized Least Square (GLS) method to generate the corresponding p-values. In this study, weight is the phenotype of interest, and the association score is calculated as the log_10(P-value). The resulting Manhattan plot was generated to visualize the significance of each SNP. To investigate the relationship between genotype and methylation we used the SciPy package pearson correlation function (scipy.stats.pearsonr) to find the correlation and p-values between SNPs and methylation sites. SNPs were filtered for sites with variance greater than 0.3 and methylation sites were filtered for sites with variance greater than 0.02, resulting in 264 methylation sites and 118 SNPs. We used the Benjamini-Hochberg procedure to find significantly correlated sites using a false discovery rate of 0.1, which identified 5 pairs as significantly correlated. The closest gene to each site in these 5 pairs were found using the UCSC genome browser (canFam4).

#### Heterozygosity

To investigate heterozygosity and runs of heterozygosity we utilized 1,656 SNP variants across 217 dogs using PLINK v1.90 (Chang et al., 2015). With the same SNP set, we estimated individual-level inbreeding coefficients (F) and runs of homozygosity (ROH) using the function detectRUNS v0.9.6 (Biscarini et al., 2018) in R v3.6.0 as:

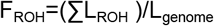

where L_ROH_ is the sum length of all ROHs detected in an individual and L_genome_ is the length of the genome analyzed. Due to the sparse density of SNPs across the genome, we used the consecutive window-free option (Marras et al., 2015) and detected tracks up to 6Mb in length with the following parameters: minSNP=10, ROHet=FALSE, maxGap=10^9^, minLengthBps=100, maxOppRun=1, maxMissRun=1.

---

Supplementary Fig 1: The coefficients assigned to each methylation site in the decision function were extracted, for each sample, by using the “coef_” attribute in scikit-learn’s logistic regression model implementation. By taking the mean absolute values of the coefficients across all samples and relabeling the sites as either being on the sex chromosome or autosome, a two-tailed t-test was performed on the coefficients in these two groups, which had a statistically significant result.

Supplementary Fig 2: The lasso model from sklearn is trained on the square root of age times the weight of dogs using leave one out cross valdiation. The x-axis is the actual square root of age times the weight, and the y-axis represents the predicted values. The correlation coefficient and the median absolute error are shown as well.

Supplementary Fig 3: A scatter plot of the two principal components that interact significantly with age in the linear model for epigenetic state produced by the Epigenetic Pacemaker (PC5 and PC7). Samples are colored by breed (only the nine most common breeds are included)

Supplementary Fig 4: Plots of genotype principal components versus weight. The blue line indicates the line of best fit using linear regression. PC7 and PC7 have the smallest p-values, both less than 0.01.

**Supplementary Figure 1:**
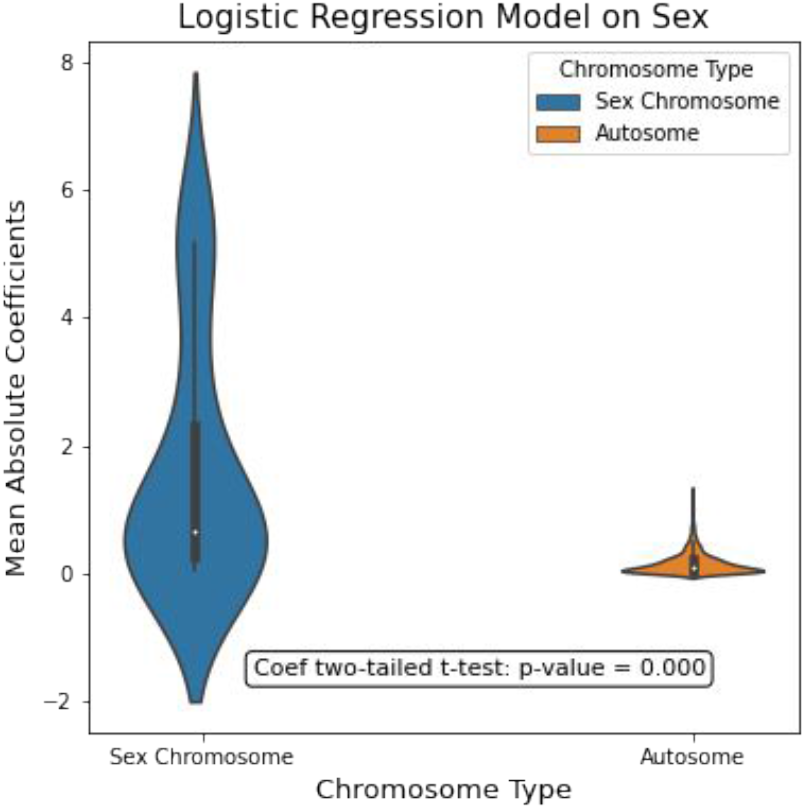
weights assigned to methylation sites in a logistic regression model for gender prediction.

**Supplementary Figure 2.**
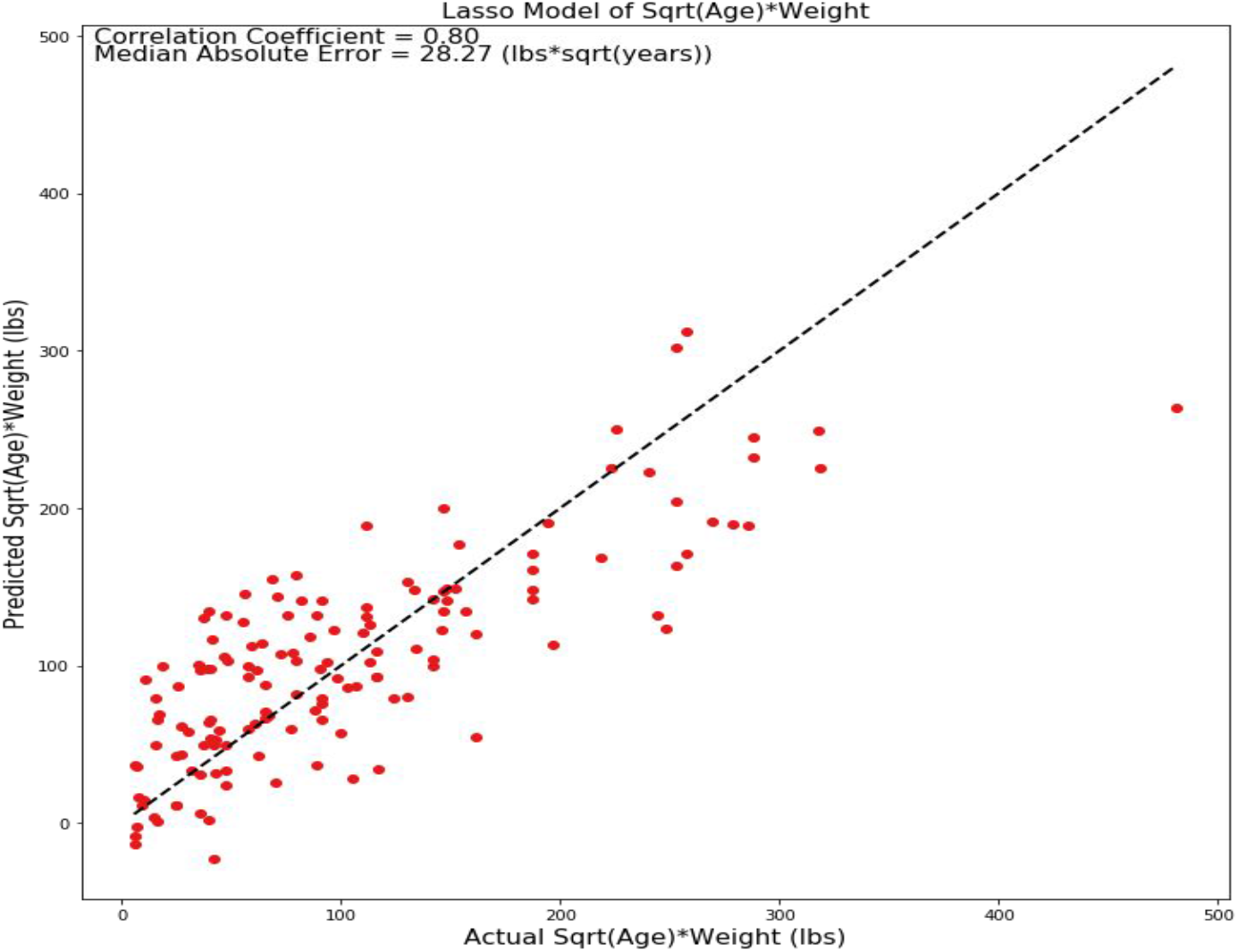
Model for square root of weight times.

**Supplementary Figure 3.**
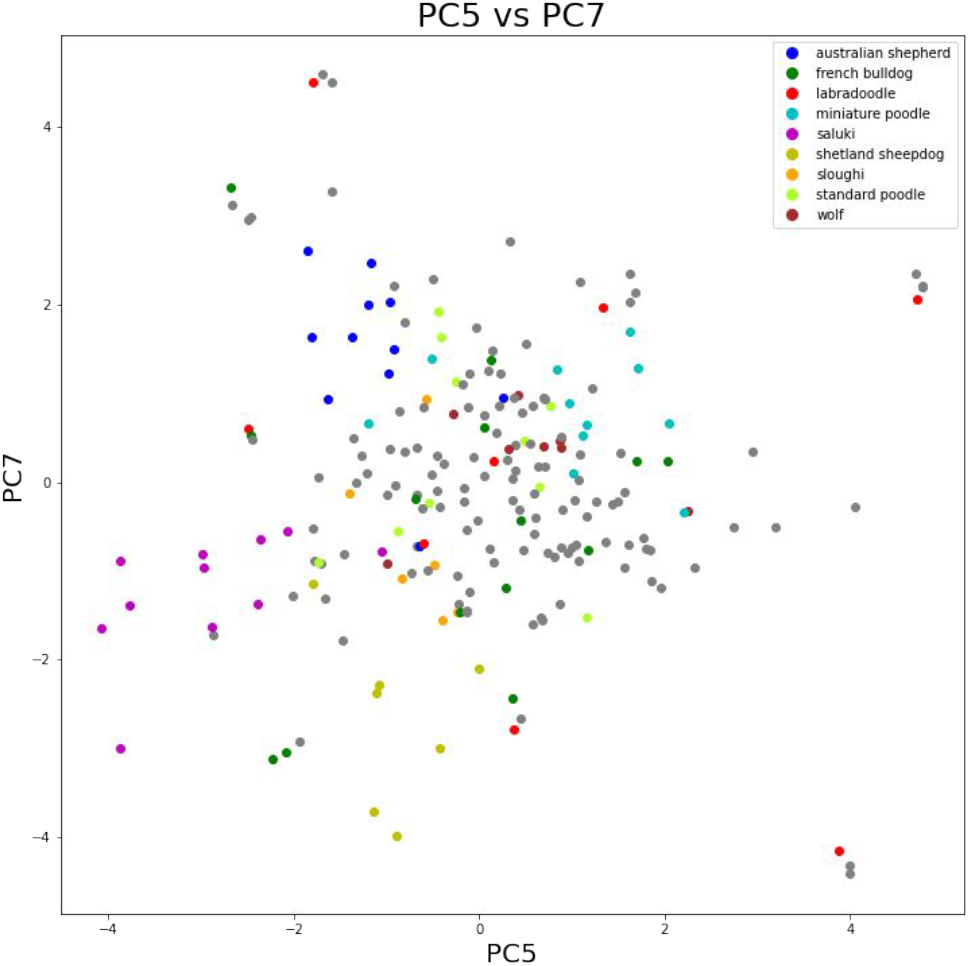
Scatter plot of PC5 vs PC 7.

**Supplementary Figure 4.**
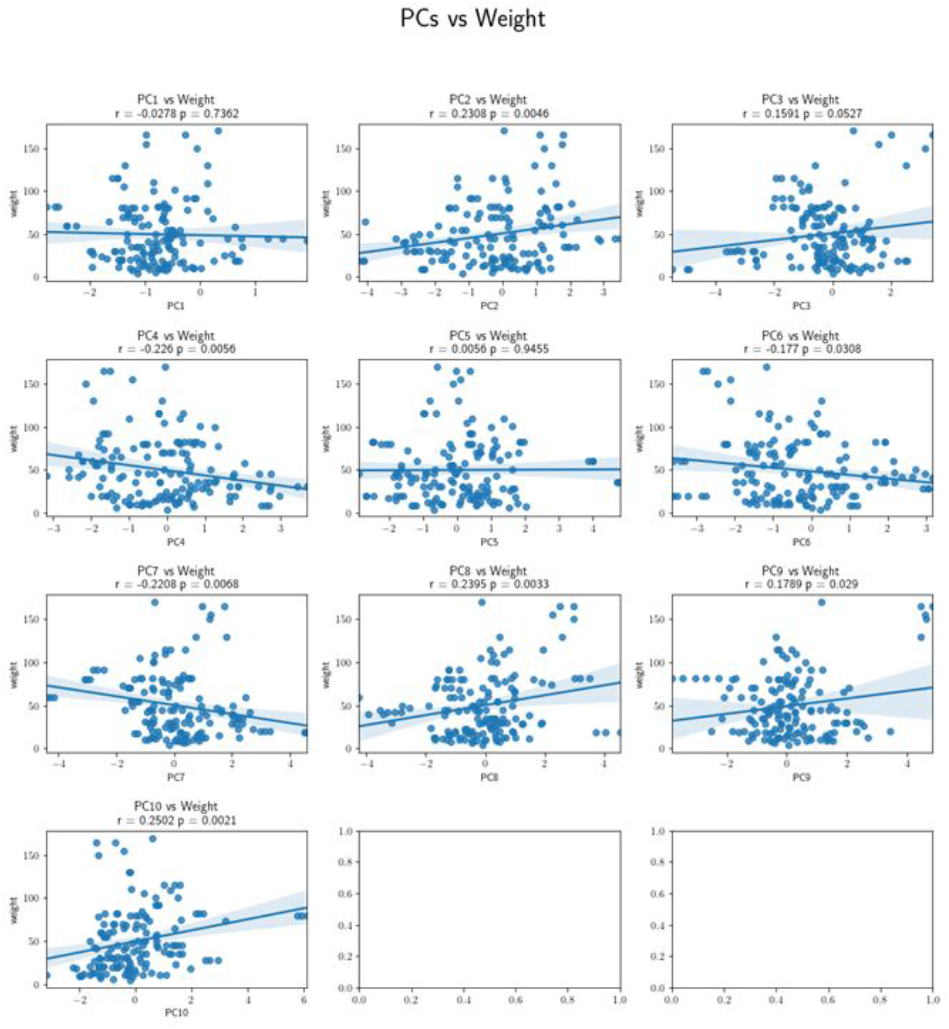
Genotype PCs vs Weight.

## Notes

### Competing Interest Statement

The authors have declared no competing interest.

## References

Almberg ES, Mech LD, Smith DW, Sheldon JW, Crabtree RL. 2009. A Serological Survey of Infectious Disease in Yellowstone National Park’s Canid Community. PLOS ONE 4:e7042. doi:10.1371/journal.pone.0007042

Biscarini F, Cozzi P, Gaspa G, Marras G. 2018. detectRUNS: Detect runs of homozygosity and runs of heterozygosity in diploid genomes. undefined.

Cassidy KA, Mech LD, MacNulty DR, Stahler DR, Smith DW. 2017. Sexually dimorphic aggression indicates male gray wolves specialize in pack defense against conspecific groups. Behavioural Processes 136:64–72. doi:10.1016/j.beproc.2017.01.011

Chang CC, Chow CC, Tellier LC, Vattikuti S, Purcell SM, Lee JJ. 2015. Second-generation PLINK: rising to the challenge of larger and richer datasets. GigaScience 4. doi:10.1186/s13742-015-0047-8

Colwell M, Drown M, Showel K, Drown C, Palowski A, Faulk C. 2018. Evolutionary conservation of DNA methylation in CpG sites within ultraconserved noncoding elements. Epigenetics 13:49–60. doi:10.1080/15592294.2017.1411447

Cubaynes S, MacNulty DR, Stahler DR, Quimby KA, Smith DW, Coulson T. 2014. Density-dependent intraspecific aggression regulates survival in northern Yellowstone wolves (Canis lupus). Journal of Animal Ecology 83:1344–1356. doi:10.1111/1365-2656.12238

Duncan CG, Grimm SA, Morgan DL, Bushel PR, Bennett BD, NISC Comparative Sequencing Program, Roberts JD, Tyson FL, Merrick BA, Wade PA. 2018. Dosage compensation and DNA methylation landscape of the X chromosome in mouse liver. Sci Rep 8:10138. doi:10.1038/s41598-018-28356-3

Faircloth BC, McCormack JE, Crawford NG, Harvey MG, Brumfield RT, Glenn TC. 2012. Ultraconserved elements anchor thousands of genetic markers spanning multiple evolutionary timescales. Syst Biol 61:717–726. doi:10.1093/sysbio/sys004

Farrell C, Snir S, Pellegrini M. 2020. The Epigenetic Pacemaker: modeling epigenetic states under an evolutionary framework. Bioinformatics 36:4662–4663. doi:10.1093/bioinformatics/btaa585

Freedman AH, Gronau I, Schweizer RM, Vecchyo DO-D, Han E, Silva PM, Galaverni M, Fan Z, Marx P, Lorente-Galdos B, Beale H, Ramirez O, Hormozdiari F, Alkan C, Vilà C, Squire K, Geffen E, Kusak J, Boyko AR, Parker HG, Lee C, Tadigotla V, Siepel A, Bustamante CD, Harkins TT, Nelson SF, Ostrander EA, Marques-Bonet T, Wayne RK, Novembre J. 2014. Genome Sequencing Highlights the Dynamic Early History of Dogs. PLOS Genetics 10:e1004016. doi:10.1371/journal.pgen.1004016

Fu K, Bonora G, Pellegrini M. 2020. Interactions between core histone marks and DNA methyltransferases predict DNA methylation patterns observed in human cells and tissues. Epigenetics 15:272–282. doi:10.1080/15592294.2019.1666649

Gilmore KM, Greer KA. 2015. Why is the dog an ideal model for aging research? Experimental Gerontology, Aging in the Wild: Insights from Free-Living and Non-Model organisms 71:14–20. doi:10.1016/j.exger.2015.08.008

Greer KA, Canterberry SC, Murphy KE. 2007. Statistical analysis regarding the effects of height and weight on life span of the domestic dog. Research in Veterinary Science 82:208–214. doi:10.1016/j.rvsc.2006.06.005

Haskins ME, Desnick RJ, Diferrante N, Jezyk PF, Patterson DF. 1984. β-Glucuronidase Deficiency in a Dog: a Model of Human Mucopolysaccharidosis VII. Pediatr Res 18:980–984. doi:10.1203/00006450-198410000-00014

Hoffman JM, Creevy KE, Promislow DEL. 2013. Reproductive Capability Is Associated with Lifespan and Cause of Death in Companion Dogs. PLOS ONE 8:e61082. doi:10.1371/journal.pone.0061082

Kang JT, Kim HJ, Oh HJ, Hong SG, Park JE, Kim MJ, Kim MK, Jang G, Kim DY, Lee BC. 2012. SRY-positive 78, XY ovotesticular disorder of sex development in a wolf cloned by nuclear transfer. Journal of Veterinary Science 13:211–213. doi:10.4142/jvs.2012.13.2.211

Kim S-A, Zhu J, Yennawar N, Eek P, Tan S. 2020. Crystal Structure of the LSD1/CoREST Histone Demethylase Bound to Its Nucleosome Substrate. Mol Cell 78:903-914.e4. doi:10.1016/j.molcel.2020.04.019

Koch IJ, Clark MM, Thompson MJ, Deere-Machemer KA, Wang J, Duarte L, Gnanadesikan GE, McCoy EL, Rubbi L, Stahler DR, Pellegrini M, Ostrander EA, Wayne RK, Sinsheimer JS, vonHoldt BM. 2016. The concerted impact of domestication and transposon insertions on methylation patterns between dogs and grey wolves. Molecular Ecology 25:1838–1855. doi:10.1111/mec.13480

Kraus C, Pavard S, Promislow DEL. 2013. The Size–Life Span Trade-Off Decomposed: Why Large Dogs Die Young. The American Naturalist 181:492–505. doi:10.1086/669665

Marras G, Gaspa G, Sorbolini S, Dimauro C, Ajmone-Marsan P, Valentini A, Williams JL, Macciotta NPP. 2015. Analysis of runs of homozygosity and their relationship with inbreeding in five cattle breeds farmed in Italy. Anim Genet 46:110–121. doi:10.1111/age.12259

Mendelson MM, Marioni RE, Joehanes R, Liu C, Hedman ÅK, Aslibekyan S, Demerath EW, Guan W, Zhi D, Yao C, Huan T, Willinger C, Chen B, Courchesne P, Multhaup M, Irvin MR, Cohain A, Schadt EE, Grove ML, Bressler J, North K, Sundström J, Gustafsson S, Shah S, McRae AF, Harris SE, Gibson J, Redmond P, Corley J, Murphy L, Starr JM, Kleinbrink E, Lipovich L, Visscher PM, Wray NR, Krauss RM, Fallin D, Feinberg A, Absher DM, Fornage M, Pankow JS, Lind L, Fox C, Ingelsson E, Arnett DK, Boerwinkle E, Liang L, Levy D, Deary IJ. 2017. Association of Body Mass Index with DNA Methylation and Gene Expression in Blood Cells and Relations to Cardiometabolic Disease: A Mendelian Randomization Approach. PLoS Med 14:e1002215. doi:10.1371/journal.pmed.1002215

Michell AR. 1999. Longevity of British breeds of dog and its relationships with sex, size, cardiovascular variables and disease. Vet Rec 145:625–629. doi:10.1136/vr.145.22.625

Morselli M, Farrell C, Rubbi L, Fehling HL, Henkhaus R, Pellegrini M. 2021. Targeted bisulfite sequencing for biomarker discovery. Methods 187:13–27. doi:10.1016/j.ymeth.2020.07.006

Parker HG, Dreger DL, Rimbault M, Davis BW, Mullen AB, Carpintero-Ramirez G, Ostrander EA. 2017. Genomic Analyses Reveal the Influence of Geographic Origin, Migration, and Hybridization on Modern Dog Breed Development. Cell Rep 19:697–708. doi:10.1016/j.celrep.2017.03.079

Pinho GM, Martin JGA, Farrell C, Haghani A, Zoller JA, Zhang J, Snir S, Pellegrini M, Wayne RK, Blumstein DT, Horvath S. 2021. Hibernation slows epigenetic aging in yellow-bellied marmots. doi:10.1101/2021.03.07.434299

Plassais J, Kim J, Davis BW, Karyadi DM, Hogan AN, Harris AC, Decker B, Parker HG, Ostrander EA. 2019. Whole genome sequencing of canids reveals genomic regions under selection and variants influencing morphology. Nat Commun 10:1489. doi:10.1038/s41467-019-09373-w

Ponder KP, Melniczek JR, Xu L, Weil MA, O’Malley TM, O’Donnell PA, Knox VW, Aguirre GD, Mazrier H, Ellinwood NM, Sleeper M, Maguire AM, Volk SW, Mango RL, Zweigle J, Wolfe JH, Haskins ME. 2002. Therapeutic neonatal hepatic gene therapy in mucopolysaccharidosis VII dogs. Proc Natl Acad Sci U S A 99:13102–13107. doi:10.1073/pnas.192353499

Poth T, Breuer W, Walter B, Hecht W, Hermanns W. 2010. Disorders of sex development in the dog—Adoption of a new nomenclature and reclassification of reported cases. Animal Reproduction Science 121:197–207. doi:10.1016/j.anireprosci.2010.04.011

Rivero S, Ceballos-Chávez M, Bhattacharya SS, Reyes JC. 2015. HMG20A is required for SNAI1-mediated epithelial to mesenchymal transition. Oncogene 34:5264–5276. doi:10.1038/onc.2014.446

Sándor S, Kubinyi E. 2019. Genetic Pathways of Aging and Their Relevance in the Dog as a Natural Model of Human Aging. Front Genet 10. doi:10.3389/fgene.2019.00948

Shah S, Bonder Marc J., Marioni RE, Zhu Z, McRae AF, Zhernakova A, Harris SE, Liewald D, Henders AK, Mendelson MM, Liu C, Joehanes R, Liang L, Heijmans BT, Hoen PAC ‘t Meurs J van, Isaacs A, Jansen R, Franke L, Boomsma DI, Pool R, Dongen J van, Hottenga JJ, Greevenbroek MMJ van, Stehouwer CDA, Kallen CJH van der, Schalkwijk CG, Wijmenga C, Zhernakova S, Tigchelaar EF, Slagboom PE, Beekman M, Deelen J, Heemst D van, Veldink JH, Berg LH van den, Duijn CM van, Hofman BA, Uitterlinden AG, Jhamai PM, Verbiest M, Suchiman HED, Verkerk M, Breggen R van der, Rooij J van, Lakenberg N, Mei H, Iterson M van, Galen M van, Bot J, Hof P van ‘t, Deelen P, Nooren I, Moed M, Vermaat M, Zhernakova DV, Luijk R, Bonder Marc Jan, Dijk F van, Arindrarto W, Kielbasa SM, Swertz MA, Zwet EW van, Levy D, Martin NG, Starr JM, Wijmenga C, Wray NR, Yang J, Montgomery GW, Franke L, Deary IJ, Visscher PM. 2015. Improving Phenotypic Prediction by Combining Genetic and Epigenetic Associations. The American Journal of Human Genetics 97:75–85. doi:10.1016/j.ajhg.2015.05.014

Shi C, Zhang M, Tong M, Yang L, Pang L, Chen L, Xu G, Chi X, Hong Q, Ni Y, Ji C, Guo X. 2015. miR-148a is Associated with Obesity and Modulates Adipocyte Differentiation of Mesenchymal Stem Cells through Wnt Signaling. Sci Rep 5:9930. doi:10.1038/srep09930

Snir S, Farrell C, Pellegrini M. 2019. Human epigenetic ageing is logarithmic with time across the entire lifespan. Epigenetics 14:912–926. doi:10.1080/15592294.2019.1623634

Stahler DR, MacNulty DR, Wayne RK, vonHoldt B, Smith DW. 2013. The adaptive value of morphological, behavioural and life-history traits in reproductive female wolves. Journal of Animal Ecology 82:222–234. doi:10.1111/j.1365-2656.2012.02039.x

Thompson EE, Nicodemus-Johnson J, Kim KW, Gern JE, Jackson DJ, Lemanske RF, Ober C. 2018. Global DNA methylation changes spanning puberty are near predicted estrogen-responsive genes and enriched for genes involved in endocrine and immune processes. Clin Epigenetics 10:62. doi:10.1186/s13148-018-0491-2

Thompson MJ, vonHoldt B, Horvath S, Pellegrini M. 2017. An epigenetic aging clock for dogs and wolves. Aging (Albany NY) 9:1055–1068. doi:10.18632/aging.101211

vonHoldt BM, DeCandia AL, Heppenheimer E, Janowitz-Koch I, Shi R, Zhou H, German CA, Brzeski KE, Cassidy KA, Stahler DR, Sinsheimer JS. 2020. Heritability of interpack aggression in a wild pedigreed population of North American grey wolves. Mol Ecol 29:1764–1775. doi:10.1111/mec.15349

vonHoldt BM, Stahler DR, Bangs EE, Smith DW, Jimenez MD, Mack CM, Niemeyer CC, Pollinger JP, Wayne RK. 2010. A novel assessment of population structure and gene flow in grey wolf populations of the Northern Rocky Mountains of the United States. Molecular Ecology 19:4412–4427. doi:10.1111/j.1365-294X.2010.04769.x

vonHoldt BM, Stahler DR, Smith DW, Earl DA, Pollinger JP, Wayne RK. 2008. The genealogy and genetic viability of reintroduced Yellowstone grey wolves. Molecular Ecology 17:252–274. doi:10.1111/j.1365-294X.2007.03468.x

Wahl S, Drong A, Lehne B, Loh M, Scott WR, Kunze S, Tsai P-C, Ried JS, Zhang W, Yang Y, Tan S, Fiorito G, Franke L, Guarrera S, Kasela S, Kriebel J, Richmond RC, Adamo M, Afzal U, Ala-Korpela M, Albetti B, Ammerpohl O, Apperley JF, Beekman M, Bertazzi PA, Black SL, Blancher C, Bonder M-J, Brosch M, Carstensen-Kirberg M, de Craen AJM, de Lusignan S, Dehghan A, Elkalaawy M, Fischer K, Franco OH, Gaunt TR, Hampe J, Hashemi M, Isaacs A, Jenkinson A, Jha S, Kato N, Krogh V, Laffan M, Meisinger C, Meitinger T, Mok ZY, Motta V, Ng HK, Nikolakopoulou Z, Nteliopoulos G, Panico S, Pervjakova N, Prokisch H, Rathmann W, Roden M, Rota F, Rozario MA, Sandling JK, Schafmayer C, Schramm K, Siebert R, Slagboom PE, Soininen P, Stolk L, Strauch K, Tai E-S, Tarantini L, Thorand B, Tigchelaar EF, Tumino R, Uitterlinden AG, van Duijn C, van Meurs JBJ, Vineis P, Wickremasinghe AR, Wijmenga C, Yang T-P, Yuan W, Zhernakova A, Batterham RL, Smith GD, Deloukas P, Heijmans BT, Herder C, Hofman A, Lindgren CM, Milani L, van der Harst P, Peters A, Illig T, Relton CL, Waldenberger M, Järvelin M-R, Bollati V, Soong R, Spector TD, Scott J, McCarthy MI, Elliott P, Bell JT, Matullo G, Gieger C, Kooner JS, Grallert H, Chambers JC. 2017. Epigenome-wide association study of body mass index, and the adverse outcomes of adiposity. Nature 541:81–86. doi:10.1038/nature20784

Wang T, Ma J, Hogan AN, Fong S, Licon K, Tsui B, Kreisberg JF, Adams PD, Carvunis A-R, Bannasch DL, Ostrander EA, Ideker T. 2020. Quantitative Translation of Dog-to-Human Aging by Conserved Remodeling of the DNA Methylome. cels 11:176-185.e6. doi:10.1016/j.cels.2020.06.006

Wang Y, Hannon E, Grant OA, Gorrie-Stone TJ, Kumari M, Mill J, Zhai X, McDonald-Maier KD, Schalkwyk LC. 2021. DNA methylation-based sex classifier to predict sex and identify sex chromosome aneuploidy. BMC Genomics 22:484. doi:10.1186/s12864-021-07675-2

Yordy J, Kraus C, Hayward JJ, White ME, Shannon LM, Creevy KE, Promislow DEL, Boyko AR. 2020. Body size, inbreeding, and lifespan in domestic dogs. Conserv Genet 21:137–148. doi:10.1007/s10592-019-01240-x

